# CCR3 antagonist impairs estradiol induced eosinophil migration to the uterus in ovariectomized mice

**DOI:** 10.1101/491860

**Authors:** Jéssica Maria Dantas Araújo, Luiz André Santos Silva, Franciel Batista Felix, Enilton Aparecido Camargo, Renata Grespan

## Abstract

The eosinophils in the uterus was associated with the successful implantation of the embryo. In addition, they are prevalent in the female reproductive tract, where they may contribute to remodeling and maintenance of epithelial integrity. However, the mechanisms by which eosinophils migrate into the uterus are still not entirely clear, since there are a variety of chemotactic factors that can cause migration of these cells. Therefore, to evaluate the CCR3 participation on eosinophil migration, we using ovariectomized C57BL/6 mice treated with CCR3 antagonist SB328437 and 17-β-estradiol (E2). Thus, it was possible to confirm the hypothesis that the CCR3 receptor has an important role in the migration of these cells to the uterus of mice, because it was clearly observed histologically that the antagonist reduces this migration and activity eosinophil peroxidase (EPO) corroborated this result. Concomitant to this, there was a 46% reduction in uterine edema, as demonstrated by the uterus weight and histology. Together, these data suggest that the CCR3 is an important receptor involved in the eosinophils regulation in uterus, including influencing the uterine edema.

## INTRODUCTION

Eosinophils are recognized for their participation in organism defense against parasites and in aggravation of asthma (1, 2). In the female reproductive system the participation of these cells has been mentioned since the 1960s (3). The first studies observed the eosinophil migration to uterus (4, 5), a phenomenon elicited by estrogen (6) but not progesterone (7).

The estrous cycle of rodents occurs within 3 to 4 days, comprising four phases, proestrus, estrus, metaestrus and diestrus (8), in response to systemic changes in sex steroids, E2. In the stages of proestrus and estrus the amount of circulating estrogen is enhance, and the rodents are susceptible to mating, thus the uterus presents a higher proportion of the granulocyte (9), possibly to ensure an effective response in the recognition of paternal antigens and / or defense against pathogens that can be transferred at the moment of mating (Mamata De, 1991). Accordingly, eosinophils performed this protective function by releasing (IL)-4 into the uterus after *Chlamydia* infection in mice (10).

During the reproductive cycle, estrogen fluctuations promote increased leukocyte turnover accompanied by tissue remodeling and a repetitive synchronization of cell proliferation, apoptosis, and differentiation (11). Thus, the administration of estrogen promotes eosinophilic migration accompanied by increase in microvascular permeability, stromal edema and proliferation of endometrial cells, besides angiogenesis, due to the secretion of vascular endothelial growth factor (VEGF) and uterine cells involved in endometrial remodeling (12, 13).

In the sense, it was postulated that interleukin (IL)-5 promotes the activation, degranulation and eosinophils migration in different tissues (14, 15), being associated with the duration of the estrous cycle (16). Furthermore, the CCR3 receptor is another important molecule for eosinophil migration (17, 18) and it is expressed in various cellular types of human endometrium and in leukocytes as eosinophil, lymphocytes and basophils (18–20). The interaction of this receptor with its ligands namely CCL (chemokine type CC binding) 11 (eotaxin 1), CCL24 (eotaxin 2) and CCL26 (eotaxin 3) produces the recruitment of eosinophils (21, 22).

Concomitant to this, several authors showed mechanisms for eosinophil migration to tissues have been identified as dependent on estrogen stimulation. Tamaki et al. associated eosinophil chemotaxis to the G protein-coupled estrogen receptor (ER) present in these cells (23). In addition, it has been shown that recruitment of estrogen-stimulated eosinophils in mammary gland depends on epidermal growth factor receptor signaling (24). More recently, Bartemes et al. have observed that type 2 innate lipoid cells (ILC2) are present in uterus in an estrogen dependent manner and highly express the ERα (25). Moreover, the ILC2s control eosinophil traffic by releasing IL-5 and local production of CCL11, in lung and adipose tissues of mice (26, 27).

The precise role of eosinophil in the homeostasis of the reproductive tract and specifically the endometrium still needs to be clarified (28). However, the literature demonstrates that eosinophils are important for the safety of embryo implantation, in addition to participating in uterine remodeling during the estrous cycle. Therefore, studies that support in understanding the mechanisms involved in migration these cells to the uterus are necessary. In this sense, we hypothesized that the CCR3 receptor effectively participates in the migration of eosinophils to the uterus of ovariectomized animals under stimulation with 17β-estradiol (E2). Our results suggest that estrogen promotes the recruitment of eosinophils into the uterus and this process is dependent on the interaction with the CCR3 receptor present in the eosinophil.

## MATERIALS AND METHODS

### Animals

All animal procedures were carried out in accordance with the standards of the Brazilian Society of Laboratory Animal Care and were approved by the Ethics Committee on the Use of Animals under protocol number 38/2014. Female C57BL/6 mice (weighing 20 to 25 g) obtained from the animal house from the Department of Physiology of the Federal University of Sergipe were used and were kept under standard housing conditions. Mice were divided into 3 experimental protocols and within them, some groups underwent surgical ovariectomy.

### Ovariectomy

The mice underwent surgical bilateral ovariectomy through a dorsal incision under anesthesia with 100 mg/kg ketamine and 10 mg/kg xylazine (Syntec^®^, Santana do Paraíba, Sao Paulo, Brazil). Additional anesthetic doses was administered throughout the experiment as necessary in order to maintain a constant level of anaesthesia, as determined from constriction of the pupil and absence of the hind limb withdrawal reflex, indicative of adequate anesthesia (29, 30). After removal of the ovaries, the dorsal wall was sutured and the cutaneous incisions closed with 10 mm clamps. As prophylaxis, animals received antiinflammatory Flunixin meglumine (EquiMed Staff^®^, Morgan Hill, CA, USA) at the dose of 2 mg/kg prior to surgery (i.m). The experimental procedures started 15 days after surgery.

### Air pouch model

The effective dose of the CCR3 antagonist in reducing eosinophil migration was determined using the air pouch model. The air pouch was induced in mice anesthetized by inhalation of halothane (Cristália^®^, Itapira, SP, Brazil). These animals were depilated in middle dorsal region, and 2.5 mL of sterile air were injected via s.c on day 0 and on day 3 the air pouch was retouch with the same volume (31). Eosinophil migration was induced on the sixth day, using recombinant mouse CCL11 (R & D System^®^, McKinley Place NE, Minneapolis, USA), reconstituted in 100 μg/mL of phosphate-buffered saline (PBS, pH 7.2) containing 1% bovine serum albumin. CCL11 (0.08 pmol/kg) was injected into the air pouch of each animal and the exudates were harvested 4 hours post-injection (32, 33). The control group received 100 μL of sterile PBS per animal.

Next, we investigated the dose of the CCR3 receptor antagonist, SB 328437 (Tocris^®^, Avonmouth, Bristol, United Kingdom) required to block eosinophil migration to the air pouch in response to CCL11. SB 328437 was dissolved in PBS and Tween 80 (0.1%) and doses of 1, 3 and 10 mg/kg were given i.p, 30 minutes before the injection of CCL11 (0.08 pmol/kg) into the air pouch (33). Furthermore one experimental group received only the stimulus with CCL11 or the stimulus PBS (negative control). Four hours after drug injection, animals were anesthetized with ketamine and xylazine and euthanized by cervical dislocation. The air pouch was washed with 1.5 mL of PBS/ethylenediaminetetraacetic acid and the lavage was collected for total and differential cell counting.

Total leukocyte number was determined in a Neubauer chamber and differential leukocyte count was performed in a cytospin preparation stained with hematoxylin- eosin for the characterization of at least 100 cells (eosinophils, neutrophils and mononuclear cells) according to the normal morphological criteria.

### Time and dose-response relationships of estrogen-induced uterine edema

To determine the time-response relationship of estrogen induced increase in uterine weight, after 15 days of ovariectomy mice received a sub-cutaneous injection of E2 (Merck^®^, Darmstadt, Germany; 100 μg/kg in the inguinal region) and 6, 12, 24 and 48 hours after the injection, the animals were anesthetized with ketamine (100 mg/kg) and xylazine (10 mg/kg) and euthanized by cervical displacement. The control group received only the vehicle, Sesame oil (S.O; Merck^®^, Darmstadt, Germany) (in volume of 60 μL). The animals had their uterus, spleen, diaphragm and bladder removed and weighed.

For the dose response experiment, the same procedure for E2 injection was followed using doses of 0.1, 1, 10 and 100 μg/kg, and organ collection was performed at the optimal time as indicated in the time-response experiments.

### Effect of SB 328437 on eosinophil migration to uterus

In order to investigate the effect of the CCR3 antagonism on eosinophil migration to uterus, a group of mice received SB 328437 (i.p) and 30 minutes later E2 (100 μg/kg) was administered; another group received vehicle solution 30 minutes before E2 and the control group received vehicle solution 30 minutes before S.O. administration. At the time indicated in the experiments to promote increased weight of the uterus, the uterus was collected, cleaned, weighed and photographed.

Evaluation of eosinophil migration was estimated by measuring eosinophil peroxidase activity analysis and cell counting in tissue sections stained with orcein as described below.

### Eosinophil peroxidase (EPO) activity

Determination of eosinophil migration to uterus was analyzed by measuring the activity of EPO using OPD (O-Phenylenediamine; Merck^®^, Darmstadt, Germany) as the substrate, according to a protocol modified from previous studies (34–36). Uterus samples (15 to 20 mg) were cut and held under ice previously to their homogenization under ice in solution (10 mmol/L, pH 6.0) containing PBS 0.005% Tween 20 (1 mL/100 mg of tissue) with a Potter homogenizer. The homogenate was centrifuged at 440 × g, 4°C for 15 minutes. The supernatant was discarded and the pellet was resuspended with the same solution and centrifuged under the same conditions. Subsequently the pellet was resuspended in PBS solution and hexadecyltrimethylammonium bromide (HTAB, 0.5%) in a ratio of 1.9 mL/100 mg of tissue and further centrifuged at 3000 × g for 15 minutes at 4°C.

The supernatant was collected and 75 μL aliquots were mixed to 150 μL of substrate prepared with Tris-HCl (0.05 mol/L, pH 8.0), OPD (1.5 mol/L) and H_2_O_2_ (6.6 mmol). After 30 minutes of incubation at room temperature, the reaction was stopped with 75 μL of H_2_SO_4_ (1 mol/L) and the absorbance determined at 492 nm on an ELISA Reader (Synergy MX^®^, Biotek, Winooski, VT, USA). In all analyzes, PBS and HTAB were used together with the substrate containing OPD, as a blank reaction. Control experiment was carried out to ensure the linearity of the detection method of EPO activity with the dilutions 1/2, 1/10 and 1/100.

### Histological analysis

After weighing the uterus and separating a fragment for eosinophil peroxidase activity measurement, the remaining tissue was immersed in 4% paraformaldehyde (for 18 h) for histological analysis. These were cut into 3 to 4 pieces and immersed in 70% alcohol until they were included in paraffin. Tissues were sliced (5 μm sections) and stained with orcein (dye solution prepared with orcein, KCN, concentrated HCl, saturated urea and 70% alcohol), specific for eosinophils (37). The morphometry of sections was performed under a microscope with a magnification of 100 x, counting all the eosinophils present in the stromal region. Data are presented as the mean of 10 microscopic fields.

### Statistical analysis

Data were expressed as mean and standard error of the mean (SEM). Statistical calculations were performed on GraphPad Prism 7.0 (Graphpad Software, San Diego, CA, USA). The specific statistical tests as well as the number of animals (n) and experimental repetitions are indicated in respective legend of the figures. *P* value <0.05 was considered as statistically significant.

## RESULTS

### Time and dose-response curve of E2-induced uterus edema

To determine the time and dose dependence of E2-induced uterine edema, the wet weight of the uterus was measured. The weight of other organs was also determined: bladder, adjacent to the reproductive tract; spleen, as a lymphoid organ; and diaphragm, representing a close muscle region of the lung. We demonstrated that after 24 hours, the E2-injected animals (100 μg/kg) presented significant uterine edema, causing increase of 68% in uterus weight compared with group that received only sesame oil (S.O; Figure 1A). Furthermore, different doses (0.1, 1, 10 and 100 μg/kg) were tested and the E2 doses of 1, 10 and 100 μg/kg were effective in inducing uterine edema 24 h after E2 administration. Nevertheless, the E2 dose of 100 μg/kg was the more effective, increasing organ weight by 67% (Figure 1B). However, this phenomenon was not observed in bladder, spleen and diaphragm weights in any time or dose evaluated (Figure 1C-H).

**Figure 1.**
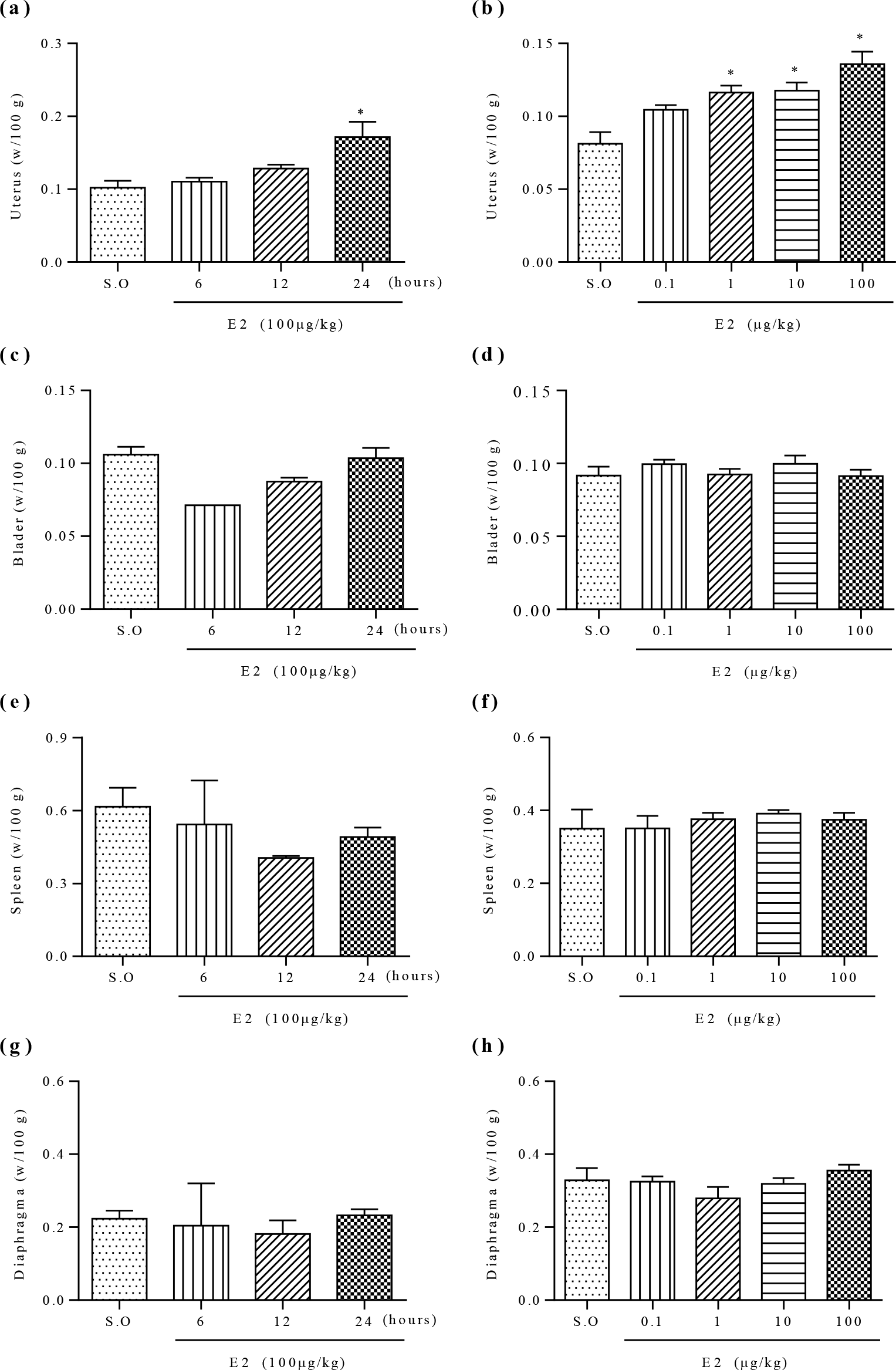
Time and dose-response curve of 17-β-estradiol (E2) in wet body weight (w). The animals were injected with E2 (100 μg/kg, s.c) and uterus (a), bladder (c), spleen (e) and diaphragm (g) were collected after 6, 12 and 24 hours of administration. The wet weight of the same organs was analyzed (b, d, f, h) after 24 hours of administration with different doses of E2 (0.1, 1, 10 and 100 μg/kg). The sesame oil group (S.O, s.c) was used as control in time of 24 hours. Data are expressed as mean and SEM of 3 independent experiments (n=5/group each). All data were analyzed using one-way ANOVAs with Bonferroni *post hoc* correction, with \**P*<0.05 when compared with S.O group.

### Evaluation of the eosinophil peroxidase (EPO) in uterus after E2 administration

Firstly, we observed that 24 hours after E2-injected animals (100 μg/kg) the EPO activity in uterine homogenates was significantly increased (60%) in uterus when compared with S.O (Figura 2A). Among the E2 doses tested (0.1 - 100 μg/kg), only the dose of 100 μg/kg increased (53%) EPO activity (Figure 2B).

**Figure 2.**
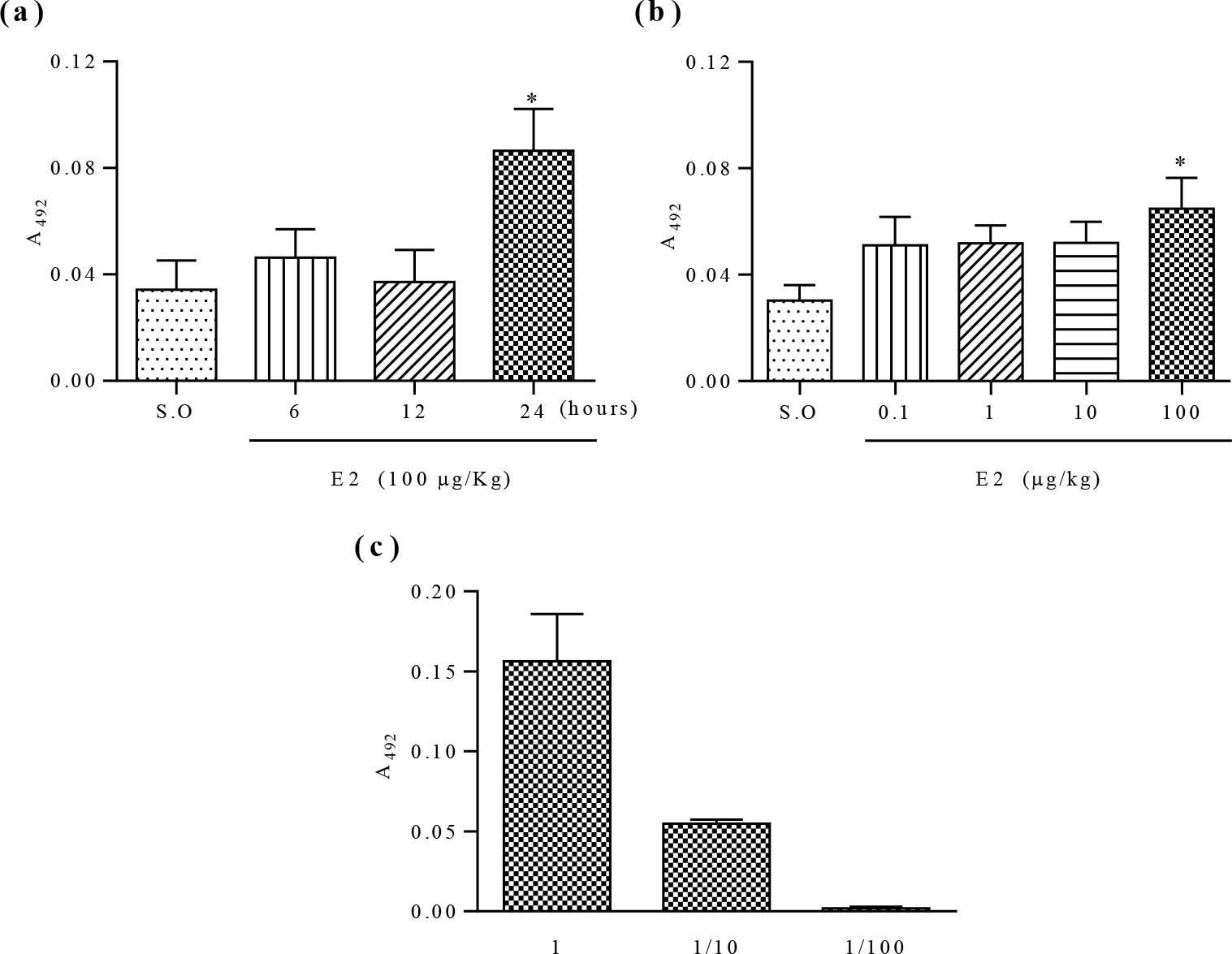
Effect of 17-β-estradiol (E2) on eosinophil peroxidase (EPO) activity in uterus. The animals were injected with E2 (100 μg/kg, s.c) and after 6, 12 and 24 hours the EPO activity was monitored as the absorbance at 492 nm. (a) (n=4 to S.O, 6, 12 and 24 h). Similarly, the assay was performed at 24-hour period with different doses of E2 (0.1, 1, 10 and 100 μg/kg) (b) (S.O n=7; 0,1 n=6; 1 n=8; 10 n=10; 100 n=10). Samples of 3 animals stimulated by E2 were diluted to demonstrate the linearity of the EPO assay (c). Animals that received only sesame oil (S.O, s.c) were used as controls. Data are expressed as mean and SEM of 2 independent experiments. All data were analyzed using one-way ANOVAs with Bonferroni *post hoc* correction, with \**P* <0.05 when compared to the S.O group.

### CCR3 antagonist (SB 328437) inhibits eosinophils migration to air pouch model

Initially, the air pouch model was used to determine the dose of the CCR3 antagonist capable of inhibiting eosinophil migration to the pouch in response to CCL11 (0.08 pmol). CCL11 increased the total leukocyte migration into the air pouch (Figure 3A), and among these leukocytes, the number of eosinophils that migrated was greater in relation to neutrophils and mononuclear cells (Figure 3B-D). In contrast, eosinophils migration to the air pouch decreased in animals treated with CCR3 antagonist at dose of 3 mg/kg or 10 mg/kg (Figure 3B). Interestingly, the CCR3 antagonist failed to affect the number of neutrophil and mononuclear cells that migrated to the pouch after CCL11 injection (Figure 3C-D).

**Figure 3.**
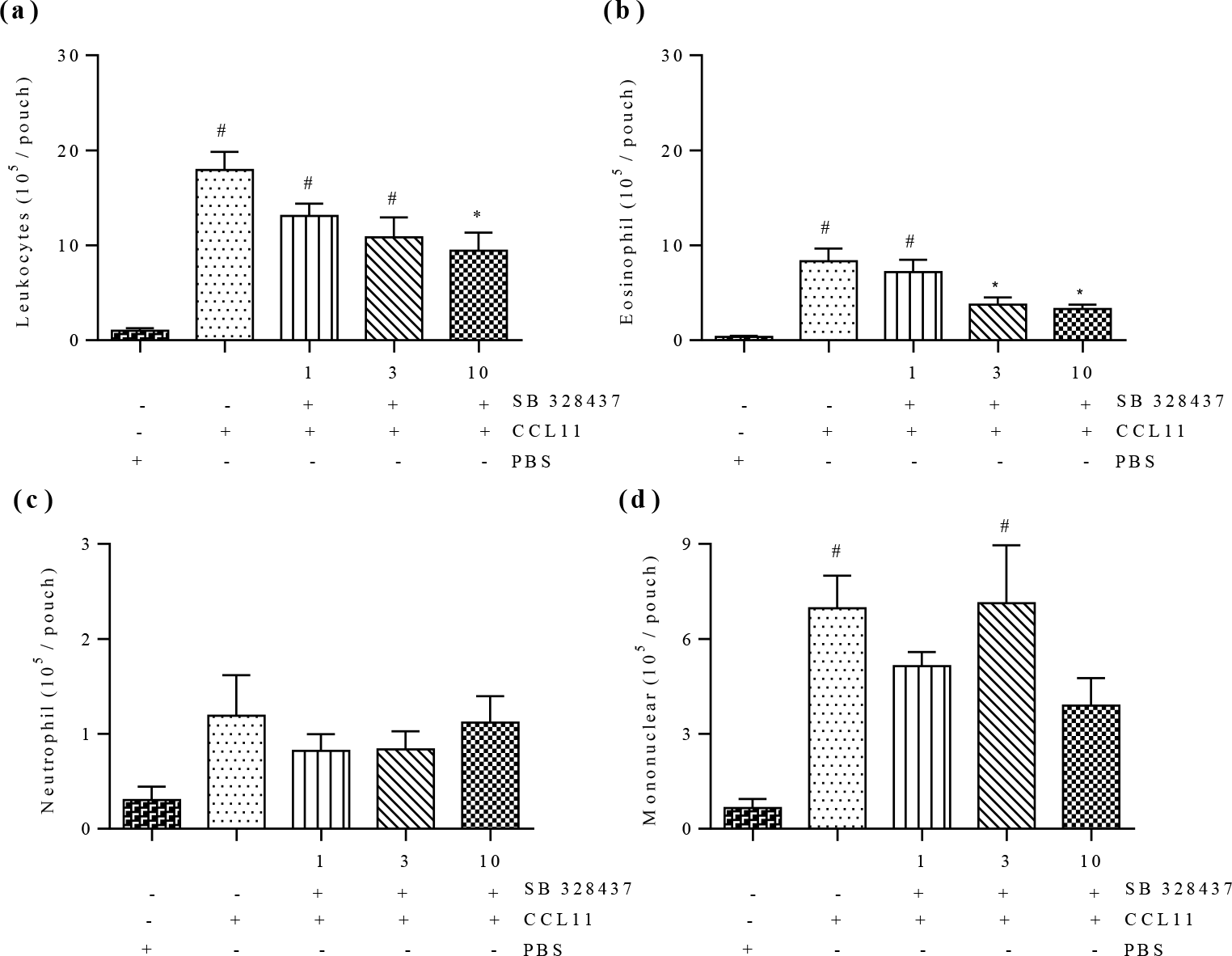
CCR3 antagonist inhibits CCL11-induced leukocyte migration to air pouch. CCL11 (0.08 pmol/kg) was administered (s.c) into the air pouch and after 4 hours, approximately 1 mL of the lavage was collected for evaluation of total leukocytes, eosinophil, neutrophil and mononuclear migration. The CCR3 antagonist was administered at the doses 1, 3 or 10 mg/kg (i.p) (n=10, n=12, n=9, respectively) 30 minutes before to administration of CCL11. After 4 hours, the washed was collected in air pouch to evaluate the migration of total leukocytes (a), eosinophils (b), neutrophils (c) and mononuclear (d). Results are expressed as mean and SEM of 2 independent experiments and all data were analyzed using one-way ANOVAs with Bonferroni *post hoc* correction, with #*P* <0.05 compared to the PBS group (negative control; n=6) and \**P* <0.05 compared to the CCL11 group (positive control; n=8).

### E2-induced uterine edema is inhibited by administration of the CCR3 antagonist

The dose of 3 mg/kg was used for the further experiments. As shown in Figure 4A, the CCR3 antagonist prevented by 46% the uterine weight gain compared to E2-injected group. In addition, there was no change in weight of bladder, spleen and diaphragm (Figure 4B-D). Corroborating this result, the macroscopic evaluation of uterus (Figure 4E) demonstrated edema (turgid appearance) and uterine hyperemia following E2 administration, when compared to S.O group, and which were greatly diminished in mice treated with the CCR3 antagonist.

**Figure 4.**
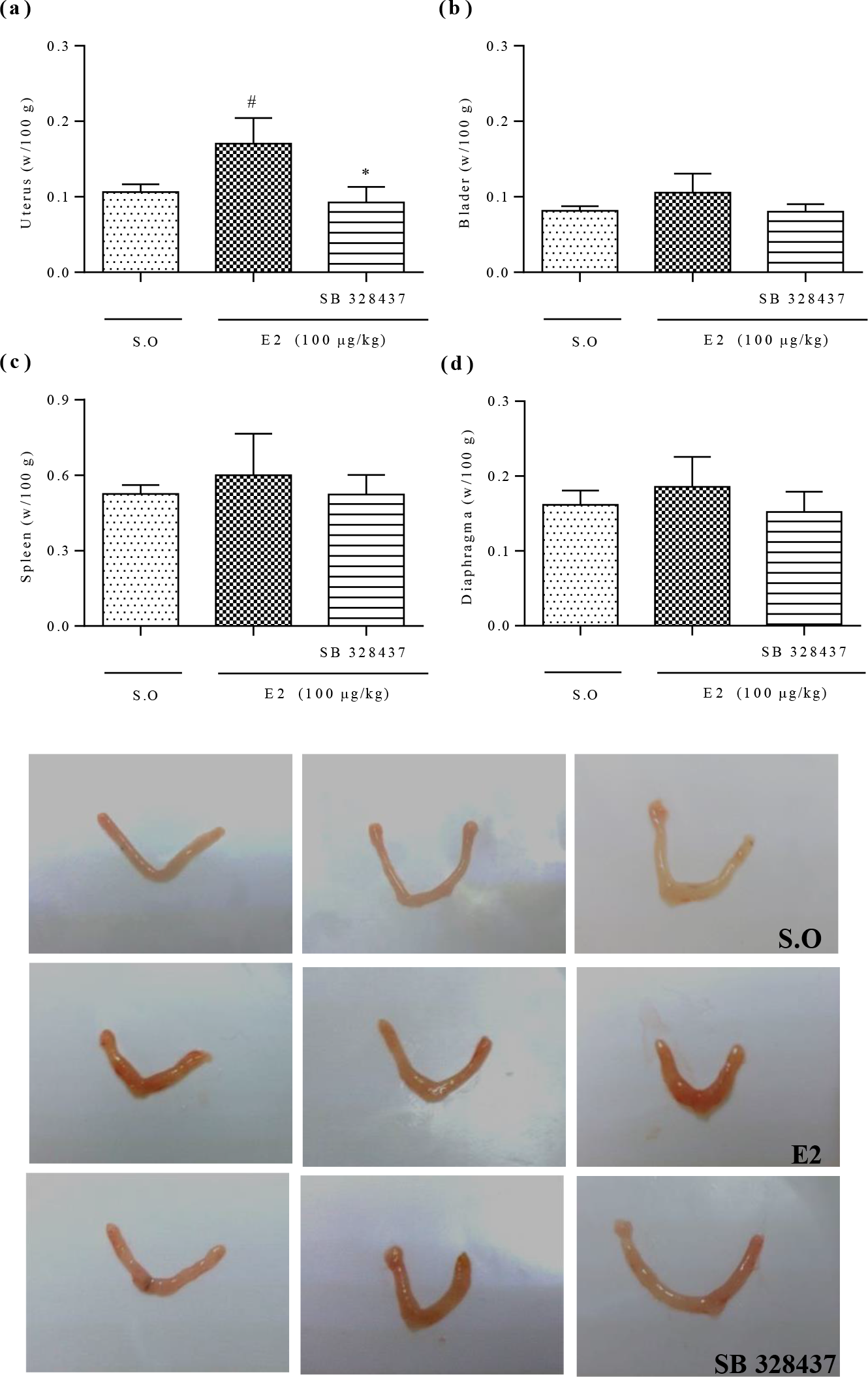
CCR3 antagonist inhibits uterus edema induced by 17-β-estradiol (E2) in mice. Groups of animals received 3.0 mg/kg of SB 328437 or vehicle (i.p; n=7) given 30 minutes before E2 injection (100 μg/kg, s.c; n=7 for the all organs) or Sesame oil (S.O, n=7). After 24 hours, the animals were euthanized and uterus (a), bladder (b), spleen (c) and diaphragm (d) were removed and weighed. In addition, representative macroscopical images of uterus are shown (e). The brightness and contrast of the figures were adjusted to better visualize the structures. Data are expressed as mean and SEM of 2 independent experiments and all data were analyzed using one-way ANOVAs with Bonferroni *post hoc* correction, with #*P* <0.05 compared to S.O group and \**P* <0.05 compared to E2 group.

### CCR3 antagonism reduces eosinophil migration E2-induced to the uterus

Eosinophil migration to the uterus was quantitated in two ways: measuring EPO activity in uterine homogenates or by direct cell counting in uterine sections stained with alkaline orcein. Figure 5A shows that CCR3 antagonist prevented the increase of EPO activity when compared to E2-injected group. In addition, uterine tissue sections stained with orcein (Figure 6) evidenced the eosinophils migration after E2 administration, clearly seen in Figure 6E and H, increasing eosinophil count by 76% compared to S.O group (Figure 6J). In contrast, panel C and G demonstrates remarkable inhibition of E2-induced eosinophil migration by the pretreatment with CCR3 antagonist, reducing 35% of the eosinophil number (Figure 6J). Also, the pretreatment with CCR3 antagonist reduced the uterine edema, as demonstrated in histology as the formation of endometrial gaps (Figure 6C and F).

**Figure 5.**
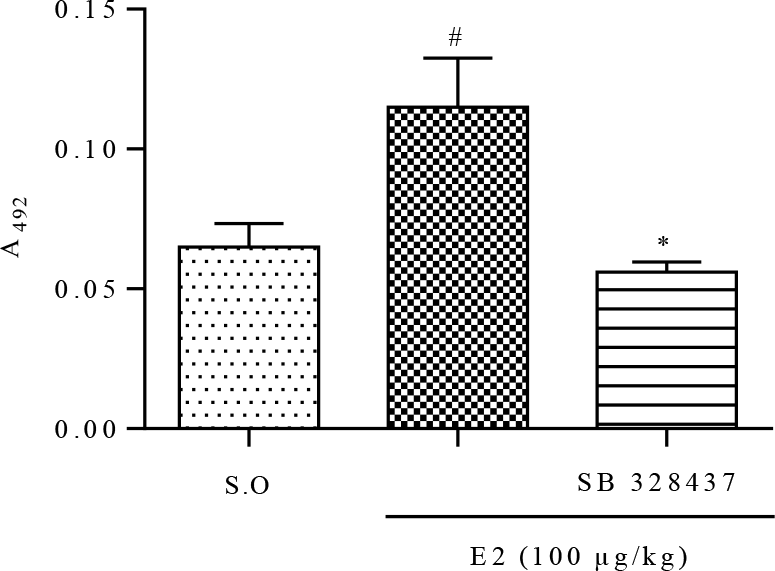
CCR3 antagonist inhibits EPO activity in uterus of E2-treated animals. Groups of animals received 3.0 mg/kg of SB 328437 or vehicle (i.p) given 30 minutes before E2 injection (100 μg/kg, s.c) or Sesame oil (S.O.). After 24 hours, the uterus was collected and EPO activity was monitored as the absorbance at 492 nm. Data are expressed as mean and SEM of 2 independent experiments and all data were analyzed using one-way ANOVAs with Bonferroni *post hoc* correction, with (n = 5/group each), with #*P* <0.05 compared to the S.O group and \**P* < 0.05 compared to the E2 group.

**Figure 6.**
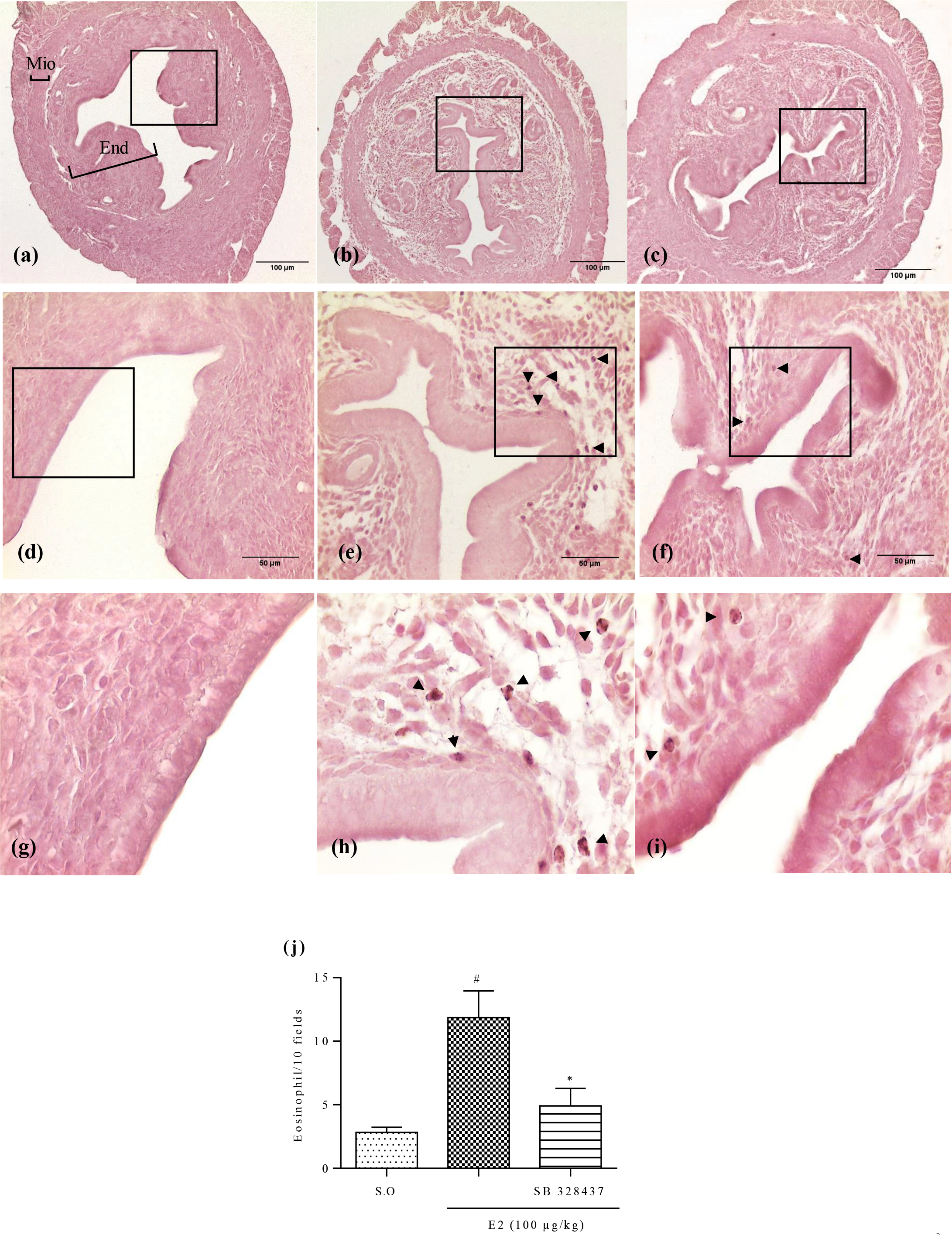
Histological analysis of uterus. Cross sections of the uterine tissue (5 μm) of ovariectomized C57BL/6 mice stained with orcein. The animals were injected with vehicle plus sesame oil (S.O) (a, d and g), vehicle plus E2 (100 μg/kg) (b, e and h) or SB 328437 (3 mg/kg) plus E2 (c, f and i) and after 24 hours uterus were collected (n = 5/group each). Panel A shows the white stripe, the uterus tissue, myometrium (Mio) and endometrium (End). Further, the regions represented by (a-c), (d-f) or (g-i) are photographs in magnification of 10, 40 or 100X, respectively. The arrows indicate eosinophils stained with orcein. The quatification of eosinophils are presented in panel (j) as the averaged count of 10 fields. The brightness and contrast of the figures were adjusted to better visualize the structures. Data are expressed as mean and SEM and were analyzed using one-way ANOVAs with Bonferroni *post hoc* correction, with #*P* <0.05 compared to S.O group and \**P* <0.05 compared to SB 328437 group.

## DISCUSSION

The findings of our study demonstrated that CCR3 antagonist was able to inhibit the eosinophils migration into the uterus as well as uterine edema of OVX mice in response to E2.

It is known that eosinophils constitutively express the CCR3 receptor in human in mice (38, 39) and the interaction with their ligands promotes recruitment of these cells to several tissues (40–42). Considering these studies cited above with results of this present work, we suggest that estrogen could be stimulating the production and/or release of CCR3 ligands by uterus resident cells. Lee et al. found in uterus of immature rats, an uterine eosinophil chemotactic factor (ECF-U) regulated by estrogen, and concluded that the estradiol-stimulated increase in uterine eosinophils is due to infiltration of these cells under the influence of ECF-U (43). Subsequently, Gouon-Evans & Pollard demonstrated that CCL11-deficient mice did not contain eosinophils in uterus and that this chemokine can be the factor ECF-U dependent of E2, found in immature rats (44). However, it is known that CCR3 is a rather promiscuous receptor, which can be activated by different ligands (45) and cause eosinophils migration into the mice uterus.

We evaluated in the air pouch model the hypothesis that CCR3-ligants whit agonist activity on the eosinophil migration could be pre-formed in the uterus. We observed that animals that received uterus extract from other E2-treated or non-treated showed higher eosinophil migration than animals that received only E2 or vehicle (sesame oil) in air pouch (data not shown). This finding supports this hypothesis since that the eosinophils migration into pouch of animals that received uterus extract without E2-treated was similar to E2- treated group.

Furthermore, Zhang et al. demonstrated that several cell types of human endometrium express the CCR3 receptor and CCL11 (46). However, other CCR3 ligands may act on eosinophilic trafficking, such as CCL5 (38), CCL7 in pulmonary granuloma (47), CCL2 (48) and CCL3 (49), in asthma. These ligands are also expressed in the uterus (50–53) and their production/release can be influenced by E2. However, CCL7 not appear to be responsible by increased migration of eosinophils in our study, because it is down-regulated by E2 (51).

It is known that, in uterus mice, cells express nuclear estrogen receptors (54) and that the interaction of estrogen with receptor α of the epithelial cells may modulate late responses after 24 hours (55). In addition, eosinophil migration occurs under E2 gene regulation (56). Concomitant to this, the stimulation by estrogen in epithelial luminal and glandular cells isolated from the human endometrium promotes CCL11 release (57). Furthermore, Moggs et al. demonstrated increased expression of the CCL11 and CCL2 gene in the mice uterus in response to the administration of E2 (58).

E2 may act directly and indirectly on gene transcription, in which the ER interacts with transcription factors in human endometrial cells (59) and this complex binds to response elements for AP-1 and NF-κB (60, 61). Thus, it is suggested that E2 acts indirectly under the transcription of the genes for CCL11 and CCL5, since these were described in mice as having the response elements for NF-κB and AP-1 (62, 63). This could cause increase on production and/or release these chemokine and eosinophils recruitment. However, this hypothesis was not substantiated by any study in mice.

We also demonstrated, for the first time, that the CCR3 antagonist reduced uterine weight in ovariectomized mice E2-treated. The E2 induced increase in uterine wet weight, since this increase is a consequence of an increase in capillary permeability our findings would indicate that CCR3 activation by uterine produced ligands in response to E2 is involved in this response. Some CCR3 antagonists have been associated with vascular endothelial growth factor (VEGF) reduction, accordingly angiogenesis (64, 65). The factor is considering major stimulator of vascular permeability and angiogenesis (66) which triggers E2-stimulated uterine edema in immature rats (67). In mice, the VEGF receptor (Flk-1/KDR) is regulated by E2 (68), and, this factor was found expressed in the secretory epithelium of the uterus, responsive to E2 (69). Thus, the relationship of CCR3 antagonism to VEGF expression or release is still not well understood. It is documented that human eosinophils release VEGF (70), which may contribute to uterine edema in response to E2 (12). If this is true for mouse eosinophils, it is possible to speculate that the amount of VEGF released by eosinophils 24 hours after single dose of E2 contributes to increased uterine weight. Robertson et al. demonstrated that gross structural changes occur in endometrium despite depletion of eosinophils, but assumed that decreased eosinophils populations may be associated with a delay in obtaining uterine responses to estrogen (71).

Moggs et al. observed a maximal uterine weight increase between 24 and 72 and VEGF expression up to 4 hours after a single dose of E2 (58). The work of Gouan-Evans previously observed increased uterine weight in mice without eosinophils, however, the animals were treated for 3 days with E2 and evaluated 24 and 48 hours after the last dose (44). Thus, it is possible that in our study the effect of VEGF on uterine weight was initiated after 24 hours of E2-treatment and was influenced by the reduction in contribution of this factor released by eosinophils.

Thereby, our study demonstrated that CCR3 is important for the eosinophils migration of into the uterus, and that the antagonism of this receptor also inhibits uterine edema 24 hours after E2-treatment. However, there are few studies in mice showing findings that support the understanding of how this modulation of E2 on CCR3- ligants.

## ACKNOWLEDGMENTS

We are mainly grateful to the intellectual contributions, throughout the process, the Prof. Dr. Gustavo Ballejo, from the Ribeirão Preto Medical School, Bazil. In addition, we appreciate for to provide perform the tissue stainig in your laboratory. His contributions were greatly appreciated and fundamental to research. We also appreciate Prof^a^. Rosilene Calazans for making the use of her histology equipment available, and assisting technically in histological procedures.

## Correspondence

Renata Grespan, Department of Physiology, Laboratory of Cellular Migration, Federal University of Sergipe, Street Marechal Rondon, no number, São Cristóvão/SE, CEP 49100-000, Brazil. telephone number +557931946643.

## Abbreviation

*CCL*: chemokine type CC binding
*CCR3*: C-C chemokine receptor type 3
*ILC2*: type 2 innate lipoid cells
*E2*: 17-β-estradiol
*EPO*: eosinophil peroxidase
*ER*: estrogen receptor
*HTAB*: hexadecyltrimethylammonium bromide
*i.p*: intraperitoneal
*IL*: interleukin
*OPD*: ortho-phenylenediamine
*PBS*: phosphate-buffered saline
*s.c*: subcutaneous
*S.O*: sesame oil
*via intramuscular*: i.m.

